# Detecting terrestrial insects from naturally exuding tree sap using environmental DNA: a pilot study

**DOI:** 10.64898/2026.05.16.724188

**Authors:** Hitoshi Kawakami, Hiroki Yuasa, Hiroki Kuroda, Tomohiro Ichinose

**Affiliations:** Graduate School of Media and Governance, Keio University, Fujisawa, Kanagawa, Japan; Faculty of Environment and Information Studies, Keio University, Fujisawa, Kanagawa, Japan

**Keywords:** environmental DNA, terrestrial eDNA, insects, tree sap, sap-mimicking trap, metabarcoding, forest monitoring

## Abstract

Terrestrial environmental DNA (eDNA) approaches are rapidly expanding, yet robust, field-ready substrates for detecting insect DNA remain limited in forest ecosystems. Tree sap is a localized microhabitat that attracts diverse insects and may provide a useful substrate for surface eDNA sampling, but its potential for insect monitoring has rarely been evaluated. Here, we present a pilot proof-of-concept study testing naturally exuding tree sap and sap-mimicking traps as terrestrial eDNA substrates. We collected swab samples from sap and trap surfaces at two forest sites in Japan (Fujisawa and Minamisanriku) and performed metabarcoding using COI and an arthropod-focused 16S marker (gInsect). Reads were processed into amplicon sequence variants and assigned by BLAST top hits against NCBI nt, with high-confidence detections defined at identity ≥98%. Across sites, sap and trap swabs yielded multiple high-confidence insect detections spanning several orders, including sap-associated stag beetles (*Dorcus* spp.). Overlap with contemporaneous conventional monitoring was limited, suggesting that sap-surface eDNA and conventional surveys capture partly different components of sap-associated insect assemblages. In a targeted 2024 spot survey, actively fermenting sap yielded multiple insect eDNA detections, whereas inactive, non-fermented sap yielded no high-confidence insect detections. Although limited by small sample size and the absence of dedicated process controls, these findings support the feasibility of tree sap as a localized terrestrial eDNA substrate and provide a basis for future replicated studies of sap-associated insect monitoring.

## 1 Introduction

Environmental DNA (eDNA) has become a powerful, non-invasive approach for detecting organisms from DNA traces in environmental samples (Bohmann et al., 2014; Taberlet et al., 2012). Early demonstrations in aquatic systems show that water eDNA can detect rare or cryptic species and characterize communities (Ficetola et al., 2008; Jerde et al., 2011). Subsequent studies have expanded this approach to community-level metabarcoding and have emphasized the importance of primer specificity and reference curation (Thomsen et al., 2012; Wilcox et al., 2013; Cristescu, 2014). In aquatic systems, eDNA metabarcoding is now widely used for biodiversity surveys and monitoring (Thomsen & Willerslev, 2015; Goldberg et al., 2016). Methodological syntheses highlight key considerations for sampling design, laboratory workflows, and bioinformatics that influence detection sensitivity (Deiner et al., 2017; Ruppert et al., 2019). Applications increasingly inform conservation and management (Lodge et al., 2012; Harper et al., 2019). eDNA has also proven useful for detecting rare species (Fukumoto et al., 2015).

However, most eDNA studies have focused on aquatic environments (Deiner et al., 2017), and terrestrial monitoring remains comparatively underdeveloped. Forest arthropods are a particularly important yet challenging target group because of their high diversity, strong seasonality, and reliance on fine-scale microhabitats (Nakamura et al., 2017; Ulyshen, 2011). Recent work shows that eDNA can also be recovered from terrestrial substrates such as forest surfaces and tree-associated materials, enabling detection of arthropods and other taxa (Allen et al., 2023a; Allen et al., 2023b). Airborne eDNA further expands terrestrial monitoring potential, although transport and deposition processes may complicate spatial inference (Johnson et al., 2021; Lynggaard et al., 2022).

Despite the rapid expansion of environmental DNA (eDNA) applications, recovering terrestrial insect eDNA remains methodologically challenging compared to aquatic systems. In terrestrial environments, DNA is highly prone to rapid degradation by UV radiation, desiccation, and microbial activity, and its dispersion is limited (Thomsen & Sigsgaard, 2019; Beng & Corlett, 2020). Recent pioneering studies have successfully recovered insect eDNA from various natural substrates, including bulk soil (Young et al., 2022), canopy leaves (Kirse et al., 2021), and airborne particles (Roger et al., 2022). However, these substrates are often highly transient, subject to rain-washing, or limited in their capacity to retain DNA over time. Therefore, identifying localized terrestrial substrates that receive frequent insect contact and can retain DNA-containing materials long enough for field sampling remains an important challenge for biodiversity monitoring.

To address this gap, we focused on tree sap as a localized and rarely evaluated substrate in forest ecosystems. Naturally exuding sap, particularly when fermenting, serves as a feeding resource and microhabitat that attracts diverse terrestrial insects, including beetles, lepidopterans, dipterans, and hymenopterans (Yoshimoto et al., 2005). Because insects repeatedly contact sap surfaces while feeding or moving, sap may retain DNA-containing biological materials such as saliva, feces, detached cells, exuviae, or body fragments. Its viscous and sticky surface may also help retain these materials locally, making sap a plausible substrate for detecting insects associated with this microhabitat. However, to our knowledge, tree sap has not yet been evaluated as a substrate for eDNA-based insect monitoring.

Reviews emphasize both opportunities and challenges for terrestrial eDNA metabarcoding, including DNA degradation, spatial heterogeneity, and reference database gaps (Hassan et al., 2022; Beng & Corlett, 2020). In this study, we evaluated whether eDNA collected from tree sap and sap-mimicking traps can yield informative species-level assignments for sap-visiting insects, and compared detections between two commonly used insect markers (COI and 16S gInsect) across two forest sites.

Metabarcoding outcomes can vary substantially with marker choice, primer-template mismatches, and the degree of DNA degradation, leading to taxon-specific biases and inconsistent detection across substrates (Elbrecht & Leese, 2015; Marquina et al., 2019). In addition, incomplete reference libraries and uneven taxonomic coverage remain major bottlenecks for species-level assignments in insects and other megadiverse groups (Weigand et al., 2019; Marques et al., 2021). To address these issues, we applied two primer sets and curated the resulting BLAST top-hit taxon lists using conservative identity thresholds and plausibility checks, then compared detected taxa with conventional sap-season monitoring records collected during the same season (2023) at each site.

Here, we conducted a pilot evaluation of sap- and trap-surface eDNA as monitoring substrates in two Japanese forest sites. Specifically, we asked: (1) whether sap and sap-mimicking trap swabs recover diverse sap-associated insect taxa to species/genus level, (2) whether COI and gInsect provide complementary detections, and (3) how high-confidence eDNA assignments (identity ≥98%) overlap with contemporaneous conventional monitoring conducted at each site in 2023.

## 2 Materials and Methods

### 2.1 Study sites and conventional monitoring

Sampling was conducted at two forest sites in Japan: Fujisawa and Minamisanriku (Fig. 2). Sample locations were recorded and labeled by substrate (sap vs sap-mimicking trap) and sample ID (Fig. 2). Conventional monitoring data used for comparison with eDNA were restricted to sap-associated insects recorded during the 2023 sap-feeding season at each site, based on visual observations and hand capture at tree sap patches. At Fujisawa, sap patches were surveyed three times per month in July and August during both daytime and nighttime (12 survey events), plus three nighttime surveys in June (15 total survey events). At Minamisanriku, surveys were conducted during a three-day campaign on 20–22 August 2023 with both daytime and nighttime searches. Although longer-term monitoring datasets are available (e.g., 2015–present at Fujisawa), we used the contemporaneous 2023 sap-season records to maximize comparability with the timing and microhabitat targeted by the eDNA swab sampling.

### 2.2 Field sampling design and substrates

We collected swab eDNA from (i) naturally exuding tree sap and (ii) sap-mimicking traps deployed on trees (Fig. 1).

**Figure 1.**
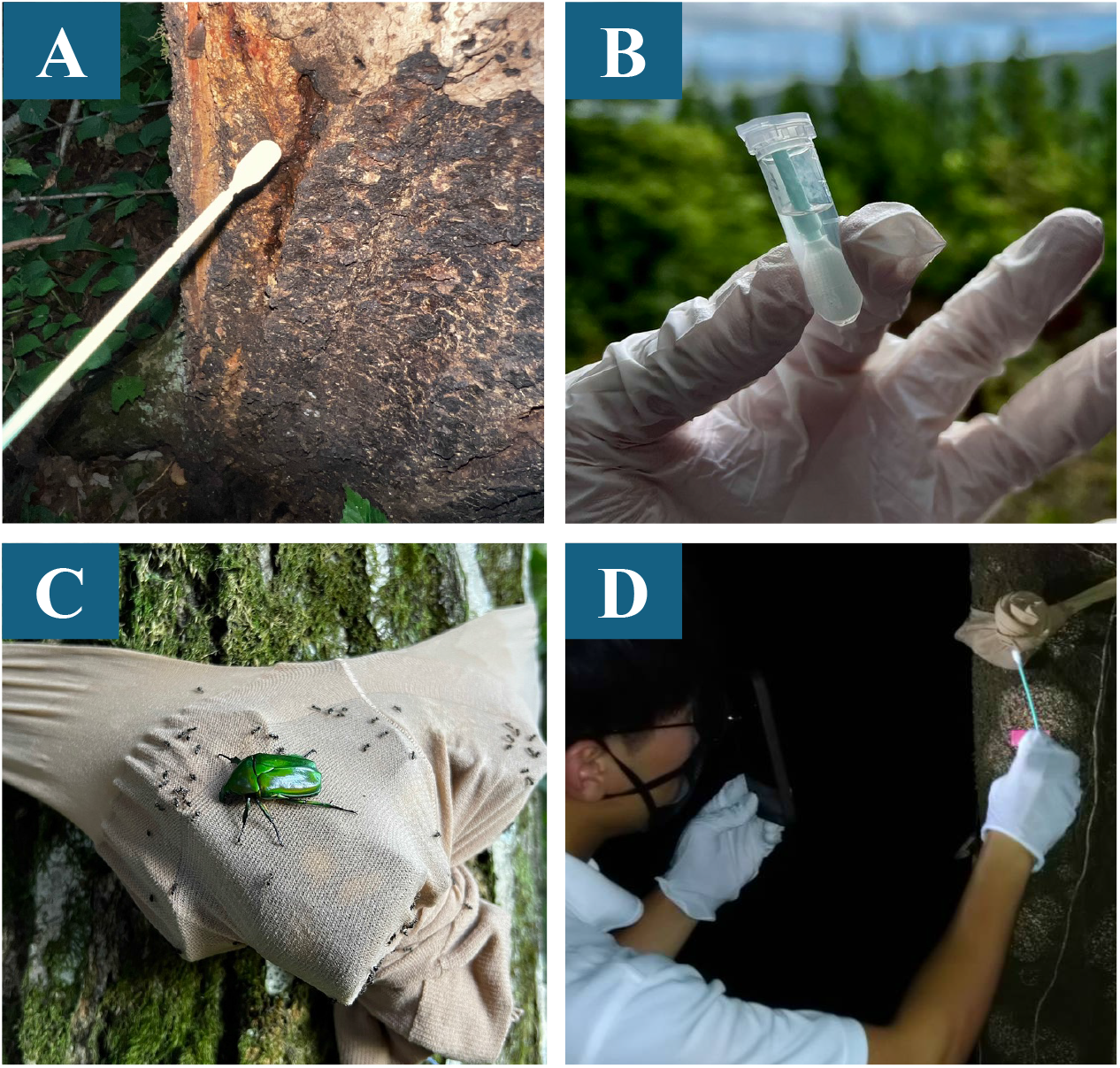
Field sampling overview for sap and sap-mimicking trap swab eDNA. (A) Swab sampling of naturally exuding tree sap. (B) Sampling swab after collection. (C) Sap-mimicking trap deployed on a tree. (D) Swab sampling of the trap surface to collect eDNA. The person shown in panel D is one of the authors of this manuscript.

Replicate samples were taken for each substrate at each site (Fig. 2).

**Figure 2.**
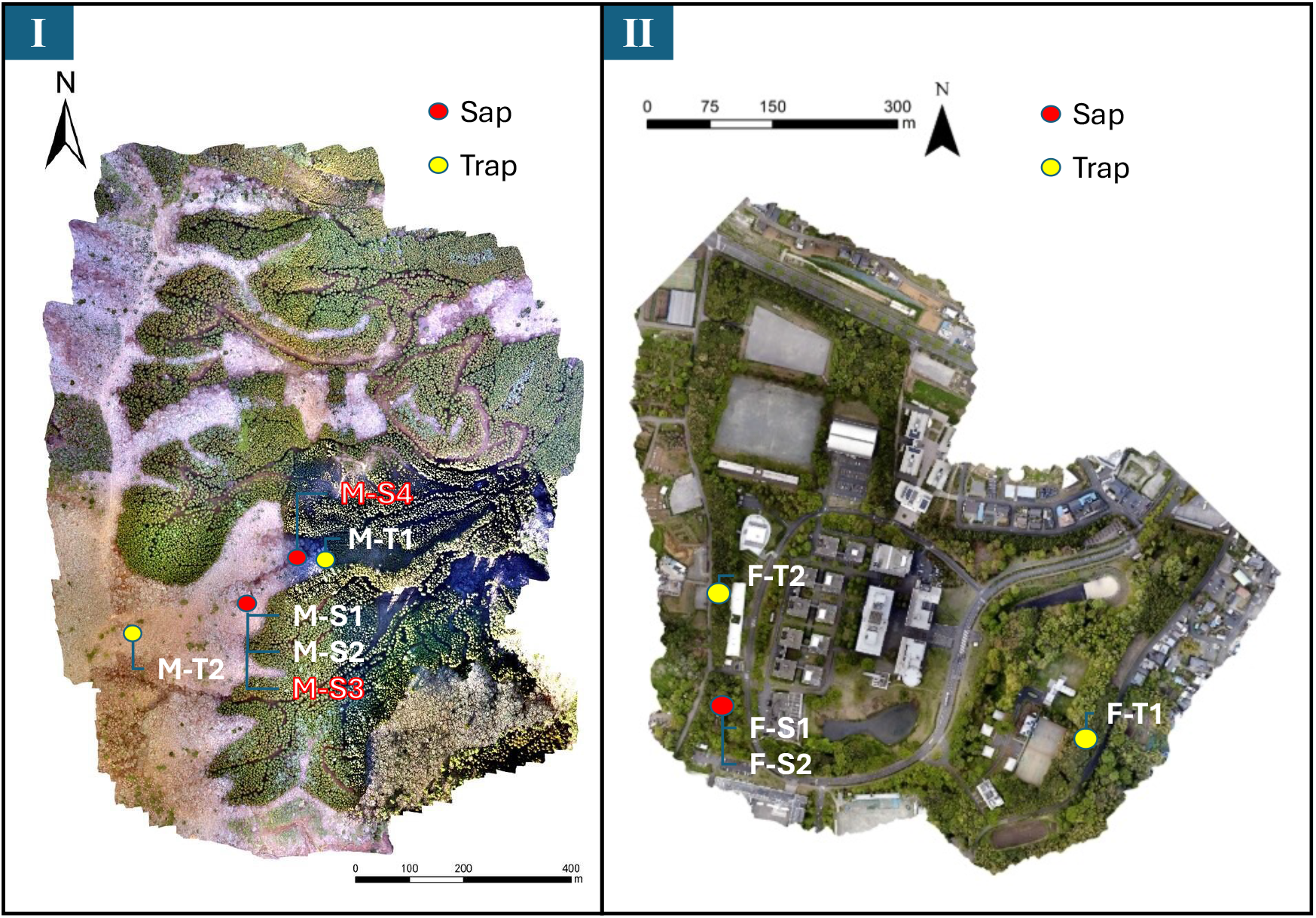
Study sites and sampling locations. Sampling was conducted at two forest sites in Japan: (I) Minamisanriku and (II) Fujisawa. Points indicate sampling locations, colored by substrate (sap vs. sap-mimicking trap). Labels show sample IDs.

### 2.3 Swab sampling procedures

#### 2.3.1 Tree sap swab sampling

For sap sampling, we selected trees with naturally exuding sap and swabbed the sap surface for ∼1 minute within a ∼10 cm × 10 cm area using sterile swabs. Two replicate samples were collected per site (four samples in total). Swabs were stored in a cooler and then frozen until analysis.

#### 2.3.2 Sap-mimicking trap swab sampling

We deployed sap-mimicking traps on trees at approximately eye level (∼1.7 m above ground). Traps were prepared using a banana fermentation bait approach: bananas were cut, mixed with shochu (∼100 mL per bunch) and dry yeast (∼3 g), fermented for ∼2 days, then packed into stockings and used as bait. Traps were attached to trees and, after ∼1 day, the trap surface was swabbed for ∼1 minute over a 10 cm × 10 cm area using the same procedure as sap swabbing. Two replicate trap samples were collected per site (four trap samples in total). Samples were cooled and then frozen until analysis.

#### 2.3.3 PBS preservation test (single sample)

As a preliminary handling test (n = 1), because cold-chain transport can be difficult in remote field conditions, one trap sample (M-T1) was preserved using diluted PBS buffer (20× diluted PBS, pH 7.4; 1 mL added) before freezing, while other samples were frozen without PBS. This PBS-preserved trap sample is annotated in Fig. 3 (Mauvisseau et al., 2021).

**Figure 3.**
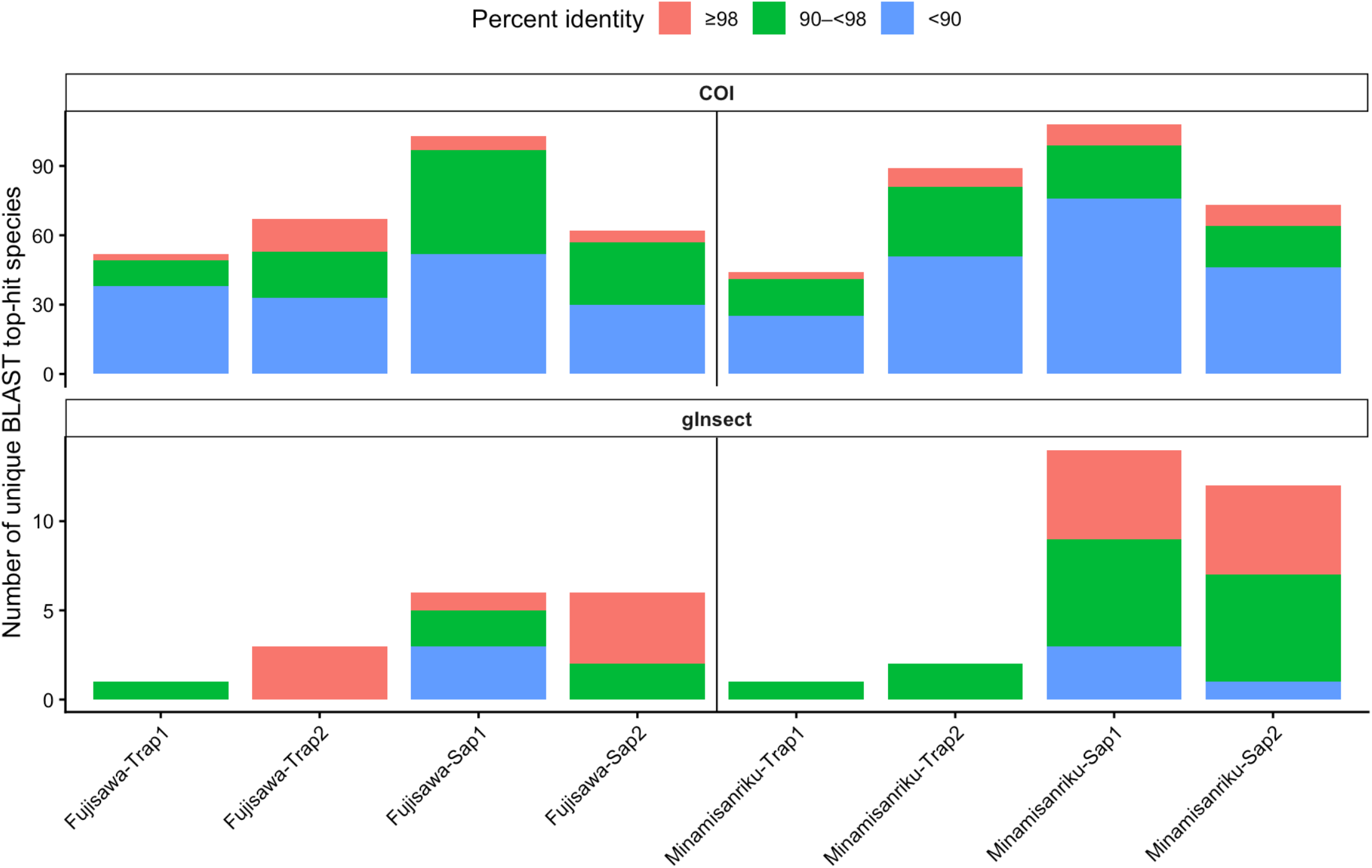
Total detections and assignment confidence from sap- and sap-mimicking trap-derived eDNA. Detected taxa were summarized as the number of unique BLAST top-hit taxa per sample and grouped by percent identity (≥98%, 90–<98%, and <90%) for (A) COI and (B) gInsect. Samples represent two sites (Fujisawa and Minamisanriku) and two substrates (sap and sap-mimicking trap). The PBS-preserved trap sample (M-T1) is annotated in the figure. This preservation comparison is unreplicated (n = 1) and should be interpreted as exploratory.

### 2.4 DNA extraction, library preparation, and sequencing

DNA extraction, library preparation, and sequencing were conducted by Bioengineering Lab. Co., Ltd. (Kanagawa, Japan). Briefly, Lysis Solution F was added to frozen samples, which were homogenized at 1,500 rpm for 2 min, incubated at 65°C for 10 min, centrifuged (12,000 × g, 2 min), and the supernatant was purified using a commercial extraction kit. DNA was quantified fluorometrically, and libraries were prepared using a 2-step tailed PCR approach. Library quality was checked using a fragment analyzer, and sequencing was performed on an Illumina MiSeq platform (2 × 300 bp; MiSeq Reagent Kit v3).

### 2.5 Metabarcoding markers

We amplified (i) a broad-spectrum mitochondrial cytochrome c oxidase subunit I (COI) marker and (ii) an arthropod-focused 16S rRNA marker (gInsect).

### 2.6 Bioinformatics and taxonomic assignment

Reads were filtered by extracting sequences matching primer starts, trimming primer sequences and 3’ terminal bases, removing low-quality reads (Q<20) and short reads (≤40 bp), and merging paired-end reads with FLASH (minimum overlap 10 bp; Magoč & Salzberg, 2011). Amplicon sequence variants (ASVs) were generated using QIIME2 (v2023.7; Bolyen et al., 2019) with DADA2 (Callahan et al., 2016) for denoising and chimera removal, followed by BLASTN searches against NCBI nt (Benson et al., 2013) (BLAST+ v2.13.0; Camacho et al., 2009; default parameters).

All analyses and visualizations were conducted in R (R Core Team, 2024).

### 2.7 Confidence thresholds and taxa curation

Detected taxa were grouped by BLAST top-hit percent identity into ≥98%, 90–<98%, and <90% bins (Fig. 3).

For comparisons with conventional monitoring, eDNA detections were defined using the identity ≥98% threshold after curation of obvious misassignments. Taxon names were standardized across datasets, and known synonyms were harmonized (e.g., *Dorcus titanus/Serrognathus platymelus*; *Dorcus striatipennis/Macrodorcas striatipennis*). Conventional records were also reduced to binomials (genus + species) to match eDNA top-hit binomials. For Fig. 4 summaries, we retained genus-level placeholders (e.g., “Genus sp.”), whereas the Venn analysis (Fig. 5) used species-level assignments only and excluded provisional “Genus sp.” labels to improve comparability with conventional records.

**Figure 4.**
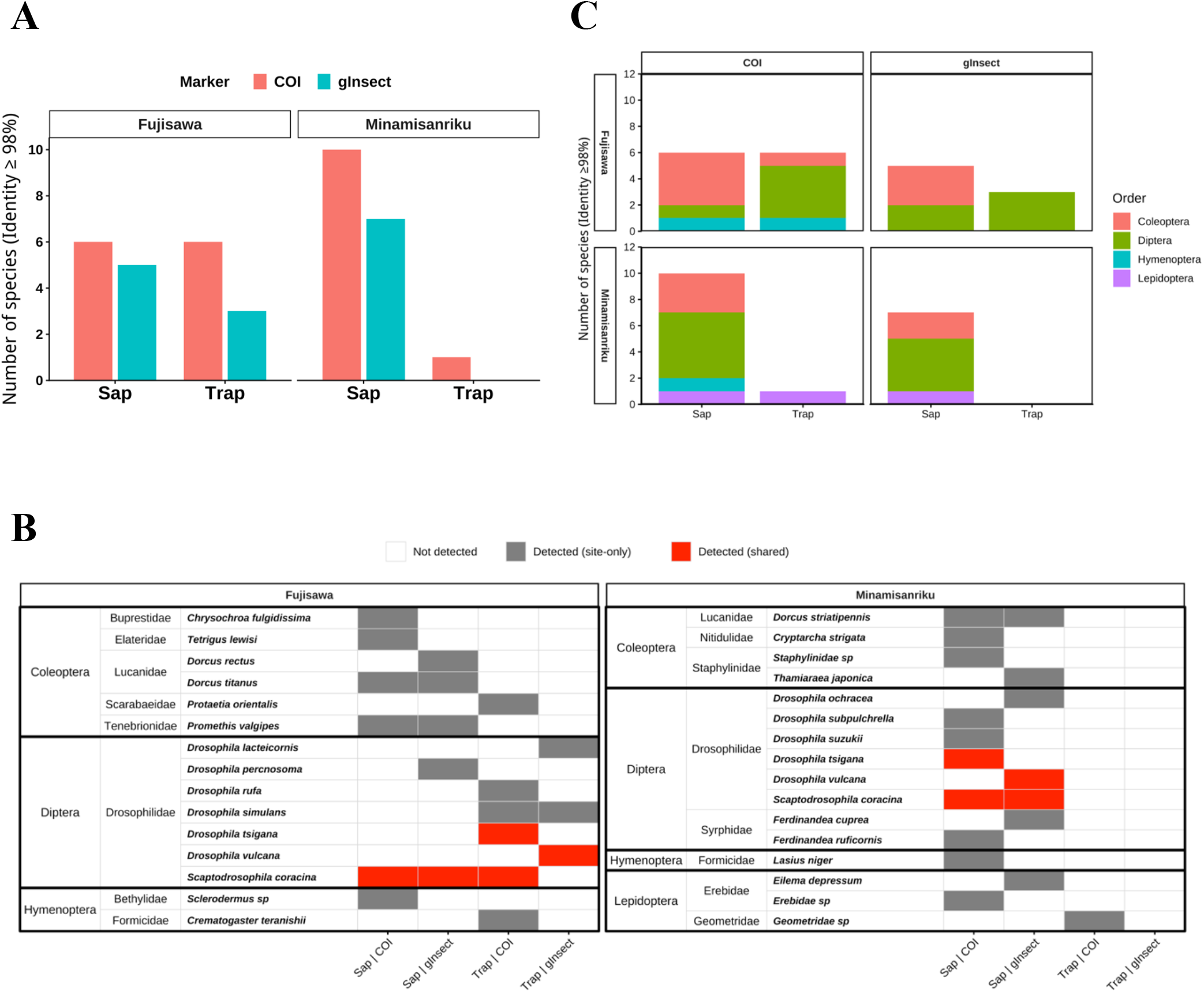
High-confidence detections (BLAST top-hit identity ≥98%) from sap and sap-mimicking trap swab eDNA across sites and markers. (A) Number of unique BLAST top-hit taxa with percent identity ≥98% detected at Fujisawa and Minamisanriku from sap and sap-mimicking trap swab eDNA using COI and gInsect. Bars show the total number of unique taxa across replicate samples within each substrate at each site. (B) Presence/absence matrix of insect taxa detected with percent identity ≥98% for each combination of substrate (sap or sap-mimicking trap) and marker (COI or gInsect) at each site. (C) Order-level composition of the detected insect taxa (identity ≥98%) for each substrate and marker at each site.

**Figure 5.**
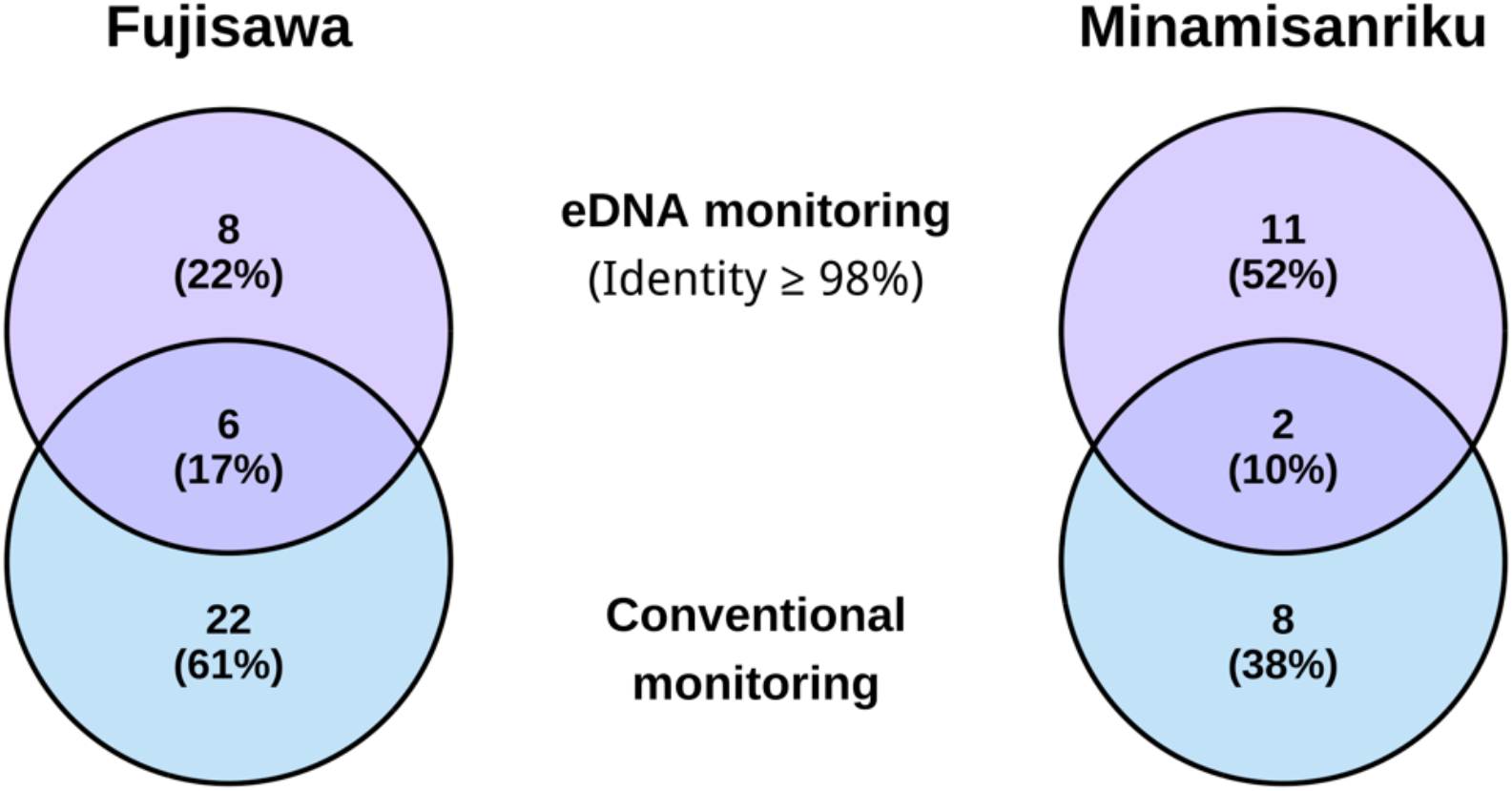
Overlap between conventional sap-season monitoring and eDNA-based monitoring at two sites. Venn diagrams show the number of insect taxa detected by conventional sap monitoring (June–August 2023 at Fujisawa; 20–22 August 2023 at Minamisanriku; conventional monitoring at sap patches) and by eDNA monitoring in Fujisawa (left) and Minamisanriku (right). For eDNA monitoring, species-level assignments were defined based on BLAST top-hit identities ≥98% after curation and synonym harmonization; provisional genus-level placeholders such as “Genus sp.” were excluded for comparability with conventional species records. Numbers indicate counts unique to each method and shared by both methods (Fujisawa: conventional only = 22, eDNA only = 8, shared = 6; Minamisanriku: conventional only = 8, eDNA only = 11, shared = 2).

### 2.8 Conventional monitoring dataset and overlap analysis

Conventional monitoring records were compiled for each site as described in Section 2.1. Because conventional monitoring and eDNA sampling differ in survey effort and detection processes, overlap values are presented descriptively and are not interpreted as direct estimates of sensitivity or false-negative rates. To compare taxonomic overlap between conventional records and eDNA detections, we constructed Venn diagrams for each site using species-level high-confidence eDNA assignments (BLAST top-hit identity ≥98%; Benson et al., 2013) after curation, synonym harmonization, and exclusion of genus-level placeholders (e.g., “Genus sp.”) as described in Section 2.7. All figures and analyses were generated in R (R Core Team, 2024).

### 2.9 Sap condition comparison in 2024

To examine whether sap condition influences insect eDNA recovery, we conducted an additional spot survey in August 2024 at Minamisanriku. Two sap swab samples were collected: MS-S3 from an “active” sap patch (fermented odor and visible insect aggregation; same patch area as MS-S1/MS-S2 sampled in 2023) and MS-S4 from a nearby tree with “inactive” sap (non-fermented liquid exudate with no visible insect activity). Sampling and laboratory processing followed the same protocols as the 2023 survey. This pilot study did not include field-blank swabs, extraction blanks, or PCR negative controls. Therefore, eDNA detections should be interpreted cautiously, particularly for taxa detected only by eDNA. The inactive sap sample MS-S4 was included as an environmental contrast sample, but it should not be regarded as a substitute for dedicated field or laboratory negative controls.

## 3 Results

### 3.1 Total detections and assignment confidence across substrates, sites, and markers

Across both sites and substrates, swab-derived eDNA yielded BLAST top-hit taxonomic assignments spanning a range of identity values. Taxa counts were summarized per sample and grouped into identity bins (≥98%, 90–<98%, <90%) for both COI and gInsect (Fig. 3). The PBS-preserved trap sample was annotated in the figure and showed different detection patterns relative to non-PBS samples; however, because this comparison was unreplicated (n = 1), it is interpreted as exploratory.

### 3.2 High-confidence detections (identity ≥98%) vary by substrate and marker

High-confidence detections (identity ≥98%) differed across sites, substrates, and markers. The number of unique high-confidence taxa (pooled across replicates) is shown in Fig. 4A, while Fig. 4B displays presence/absence patterns by substrate × marker at each site. Order-level composition of detected insect taxa is summarized in Fig. 4C.

Across both forest sites, sap and sap-mimicking trap swab eDNA yielded multiple high-confidence insect detections (BLAST top-hit identity ≥98%), spanning several orders, including Diptera, Coleoptera, Lepidoptera, and Hymenoptera. At Fujisawa, sap swabs yielded 6 COI taxa and 5 gInsect taxa, while sap-mimicking trap swabs yielded 6 COI taxa and 3 gInsect taxa (Fig. 4A). At Minamisanriku, sap swabs yielded 10 COI taxa and 7 gInsect taxa, while sap-mimicking trap swabs yielded 1 COI taxon and no gInsect taxa (Fig. 4A). Summed across substrates and markers, we detected 15 insect taxa at Fujisawa and 16 insect taxa at Minamisanriku (Fig. 4B), including several provisional assignments (e.g., “Genus sp.”). Excluding provisional genus-level assignments, species-level detections totaled 14 taxa at Fujisawa and 13 taxa at Minamisanriku (used for the Venn comparisons in Fig. 5). Marker-specific patterns were evident: some taxa were detected by both COI and gInsect, while others were recovered predominantly by a single marker, underscoring complementarity between primer sets for broad arthropod surveillance.

### 3.3 Limited overlap between conventional monitoring and eDNA monitoring

Overlap between conventional monitoring and eDNA monitoring (identity ≥98%, curated as described above) was limited at both sites (Fig. 5). In Fujisawa, conventional monitoring detected 22 taxa not detected by eDNA, eDNA detected 8 taxa not recorded by conventional monitoring, and 6 taxa were shared. In Minamisanriku, conventional monitoring detected 8 taxa not detected by eDNA, eDNA detected 11 taxa not recorded by conventional monitoring, and 2 taxa were shared. Shared taxa included *Dorcus titanus* at Fujisawa and *Dorcus striatipennis* and *Lasius niger* at Minamisanriku.

### 3.4 Sap condition was associated with eDNA detection in the 2024 spot survey

Sap condition was associated with differences in insect eDNA detection in the 2024 spot survey at Minamisanriku (Fig. 6). The active/fermented sap sample (MS-S3) yielded 8 high-confidence insect taxa with COI and 8 taxa with gInsect, including overlap with the 2023 sap samples (MS-S1/MS-S2) of 4 taxa (COI) and 3 taxa (gInsect). In contrast, the inactive/non-fermented sap sample (MS-S4) yielded 0 taxa with COI and 0 taxa with gInsect at the identity ≥98% threshold. These observations suggest that visible insect activity and fermentation status—likely linked to visitor contact rates—may influence recoverable surface eDNA signals.

**Figure 6.**
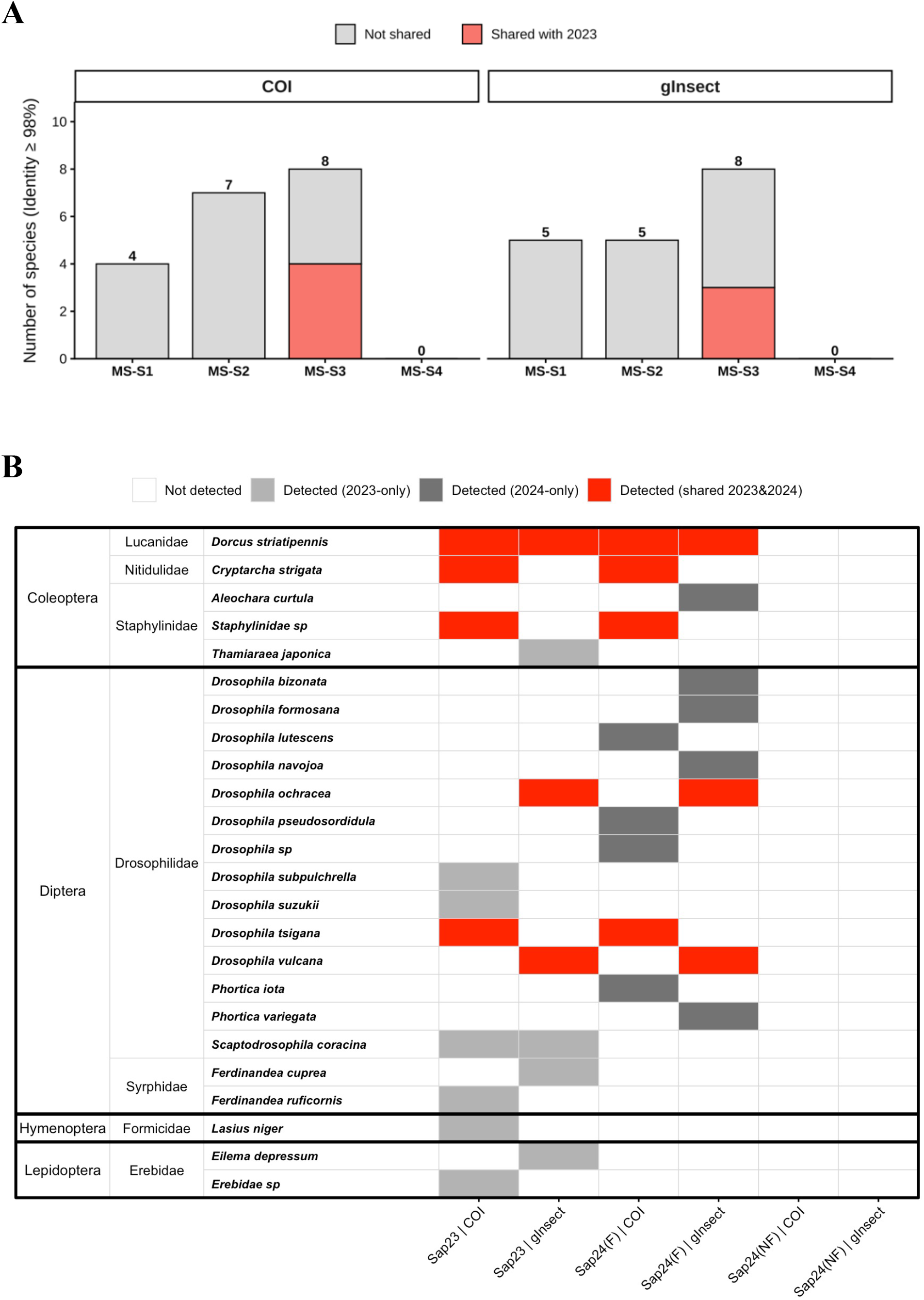
Influence of sap condition on insect eDNA detections at Minamisanriku (2024 spot survey). (A) High-confidence insect taxa (BLAST top-hit identity ≥98%) detected from MS-S3 (active/fermented sap with visible insect activity) and MS-S4 (inactive/non-fermented sap with no visible activity) using COI and gInsect. (B) Overlap of detected taxa between 2023 sap samples (MS-S1/MS-S2) and the 2024 samples (MS-S3/MS-S4).

**Figure 7.**
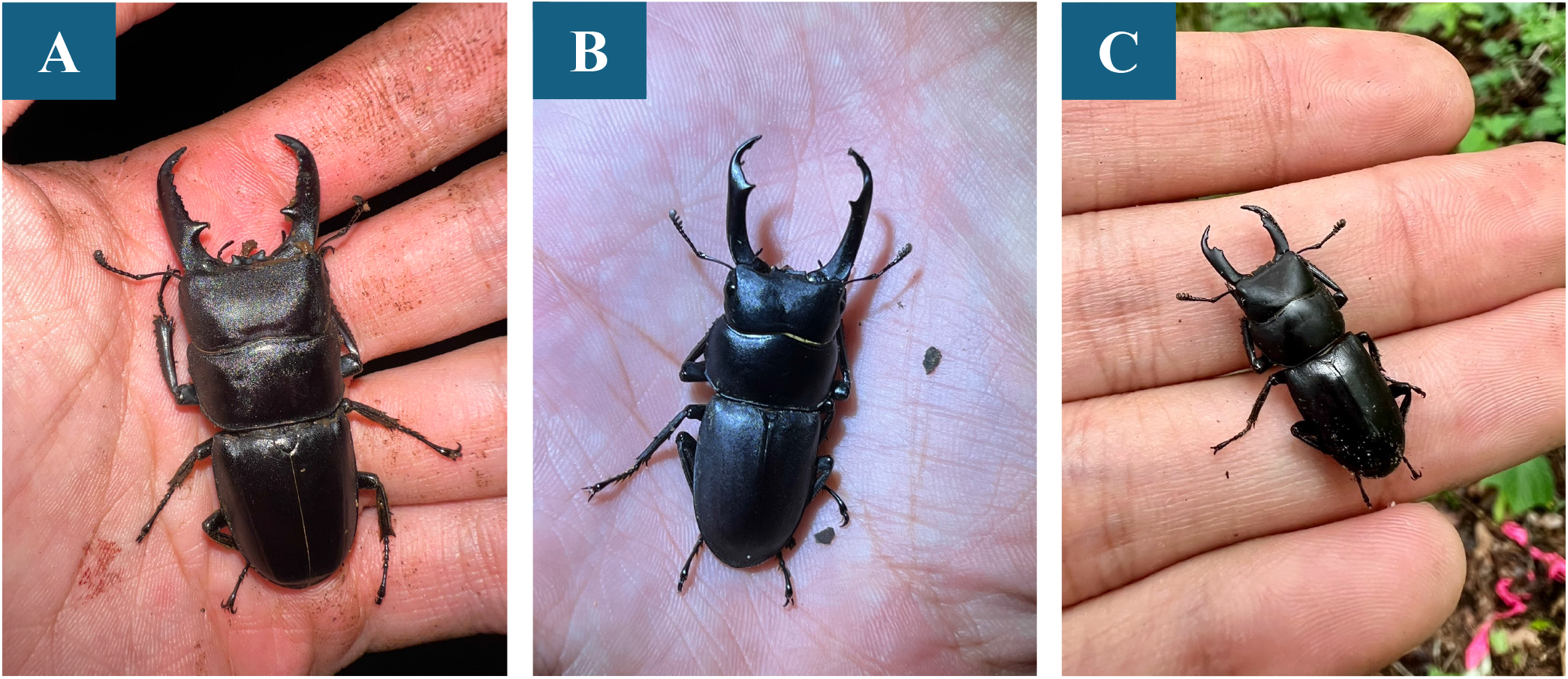
Representative sap-associated stag beetles detected by both conventional sap-season monitoring and sap eDNA in 2023. (A) *Dorcus titanus* from Fujisawa (listed as Vulnerable (VU) in Kanagawa Prefecture), (B) *Dorcus rectus* from Fujisawa, and (C) *Dorcus striatipennis* from Minamisanriku. Conservation status was checked using the Ministry of the Environment ‘Ikimono Log’ prefectural threatened species search (Ministry of the Environment, Japan, n.d.; retrieved 19 February 2026).

## 4 Discussion

### 4.1 Main findings

Our swab-based terrestrial eDNA approach recovered a taxonomically diverse set of sap-associated insects from both natural tree sap and sap-mimicking traps at Fujisawa and Minamisanriku. High-confidence taxon assignments (BLAST top-hit identity ≥98%) included multiple orders (Diptera, Coleoptera, Lepidoptera, and Hymenoptera), indicating that surface eDNA collected from small forest microhabitats may capture informative signals of arthropod diversity. This aligns with recent terrestrial surface eDNA studies showing that DNA left on vegetation or other contacted surfaces can be informative for arthropod community assessment (Allen et al., 2023b; Macher et al., 2023; Yoneya et al., 2023). We also detected insects not recorded by sap-season visual surveys at each site, suggesting that sap and trap swab eDNA may complement conventional monitoring by providing additional detections of cryptic, nocturnal, or rarely observed taxa (Bohmann et al., 2014; Ruppert et al., 2019).

### 4.2 Complementarity of markers

Comparisons across primer sets indicated clear marker-specific detection patterns: some taxa were detected by both COI and gInsect, while others were recovered only by one primer set. Such complementarity is expected for arthropod metabarcoding because primer binding sites, amplicon length, and taxonomic coverage differ among markers, leading to differential amplification efficiencies across taxa (Elbrecht & Leese, 2017; Porter & Hajibabaei, 2018). Benchmarking studies and reviews also recommend multi-marker approaches when broad community coverage is needed, particularly in diverse arthropod assemblages (Elbrecht et al., 2019; Weigand et al., 2019; Zinger et al., 2019).

### 4.3 Relationship to conventional monitoring records

Overlap between eDNA detections and conventional sap-season records was limited (Fig. 5). Because our comparison used sap-associated insects recorded during the same season (2023), this discrepancy likely reflects differences in detection processes rather than a simple temporal-scale artifact. Conventional monitoring records the presence of visible individuals during survey periods, while sap eDNA may integrate DNA left by brief or nocturnal visits and may capture taxa that are difficult to observe or identify in the field (Montgomery et al., 2021; Ruppert et al., 2019). The larger number of conventional-only taxa at Fujisawa may also reflect greater conventional survey effort at this site, including repeated daytime and nighttime surveys across June–August. Therefore, overlap values should not be interpreted as direct estimates of detection sensitivity or false-negative rates. Instead, they provide a descriptive comparison between two methods that differ in sampling effort, temporal integration, and detection processes.

The eDNA-only detections (8 taxa in Fujisawa and 11 taxa in Minamisanriku; Fig. 5) suggest that sap-surface eDNA may complement conventional surveys by detecting taxa that were not observed during visual monitoring. Some of these detections involved small or cryptic taxa that may be easily overlooked in the field, such as the drosophilid fly *Scaptodrosophila coracina* and the nitidulid beetle *Cryptarcha strigata*. In addition, primer bias, variation in DNA shedding, stochastic detection of low-template DNA, and imperfect reference databases can reduce congruence between methods even when sampling the same microhabitat (Elbrecht & Leese, 2015; Weigand et al., 2019). Despite these limitations, shared sap-associated taxa were detected at both sites (Fujisawa: 6 taxa; Minamisanriku: 2 taxa), supporting the interpretation that sap-targeted eDNA may serve as a complementary tool to conventional sap surveys.

Representative taxa further illustrate the potential complementarity between sap-surface eDNA and conventional monitoring. At Fujisawa, *Dorcus titanus* was detected by both methods. This species is listed as Vulnerable (VU) in Kanagawa Prefecture, and its nocturnal and cavity-dwelling behavior can make visual detection difficult; in 2023, it was observed during only one of 15 conventional survey events despite targeted effort. In contrast, *Chrysochroa fulgidissima* was detected only by eDNA and was not recorded during the 2023 sap-season conventional surveys. Because sap visitation by this species was not visually confirmed during our surveys, this detection should be interpreted cautiously and may reflect brief or infrequent sap visitation, indirect DNA deposition, or low-level contamination. These examples suggest that sap-surface eDNA may complement conventional monitoring by detecting rarely observed taxa and by generating hypotheses about microhabitat use that can be tested with targeted observations.

### 4.4 Methodological considerations and limitations

Although we restricted inference to high-identity BLAST top hits, reference database incompleteness remains a key limitation for terrestrial arthropod eDNA, particularly for regional faunas with many undescribed or poorly represented taxa (Ratnasingham & Hebert, 2007; Weigand et al., 2019). Even large general repositories and metabarcoding datasets exhibit uneven taxonomic coverage and annotation issues that can propagate into species assignments (Benson et al., 2013; Shokralla et al., 2014; Zinger et al., 2019). This may explain the occurrence of multiple ‘sp.’ assignments and taxa lacking Japanese common names. Another important consideration is the use of a top-hit identity threshold to define species detections. While a high threshold reduces false positives, it can also increase false negatives for taxa with limited reference sequences or high intraspecific variation. Future work should evaluate alternative assignment strategies (e.g., lowest common ancestor approaches or curated local reference libraries) and sensitivity analyses across identity thresholds to quantify robustness (Zinger et al., 2019).

Surface eDNA signals may also be influenced by DNA persistence, transport, and shedding rates (Andruszkiewicz Allan et al., 2021). Sap is a viscous substrate that may protect DNA from degradation and allow accumulation across multiple visits. In contrast, trap surfaces and exposed sap patches may differ in UV exposure, rainfall, and microbial activity, all of which can affect persistence and therefore comparability across substrates and sites (Barnes & Turner, 2016; Deiner et al., 2017). Rigorous field and laboratory controls are therefore essential, and repeated temporal sampling will help distinguish persistent background DNA from recent visitation signals. Finally, because sap-associated communities are microhabitat-specific, eDNA from sap should be interpreted as an indicator of sap-visiting assemblages rather than a complete representation of the surrounding forest insect community.

Nevertheless, although contamination cannot be fully excluded, the detection of ecologically plausible sap-associated taxa, such as *Dorcus* spp. and *Cryptarcha strigata*, supports the biological plausibility of the eDNA signals recovered.

### 4.5 Implications and future directions

To our knowledge, this study is among the first to report species- and genus-level identification of sap-visiting insects from terrestrial eDNA collected directly from tree sap and sap-mimicking traps. Terrestrial eDNA metabarcoding is rapidly developing, but many applications still focus on soil, litter, or general surface swabs rather than localized ‘hotspot’ substrates such as tree sap (Hassan et al., 2022; Beng & Corlett, 2020). Recent studies indicate that forest surfaces and tree-associated substrates can capture informative eDNA signals for arthropods and other taxa, supporting the feasibility of non-water-based monitoring in forest ecosystems (Allen et al., 2023a; Allen et al., 2023b).

Taken together, we position these findings primarily as a proof of concept: tree sap and sap-mimicking traps can yield informative terrestrial eDNA signals, but broader ecological inference will require increased replication, seasonal coverage, and more standardized sampling protocols.

### 4.6 Substrate selection and insect-active sap patches

Sap-mimicking traps may provide a deployable and more standardized sampling unit, but they should not be assumed to be equivalent to naturally active sap patches. In this pilot dataset, trap-derived detections differed among sites and were particularly limited at Minamisanriku, suggesting that bait composition, fermentation stage, deployment duration, microclimate, and insect visitation rates may influence eDNA recovery. Future work should optimize trap design and directly compare trap surfaces with naturally active sap patches under replicated conditions.

The contrasting 2024 sap samples indicate that tree sap is not a uniform eDNA substrate. An actively fermenting sap patch with visible insect aggregation (MS-S3) yielded multiple high-confidence detections, while a visually inactive, non-fermented sap exudate (MS-S4) yielded none at the identity ≥98% threshold. This contrast suggests that sap-surface eDNA is most informative when collected from insect-active sap patches, supporting its interpretation as a microhabitat- and activity-dependent indicator of sap-visiting assemblages rather than a comprehensive census of local insect diversity. However, because this comparison was unreplicated and lacked dedicated process controls, the result should be interpreted as preliminary. Future studies should test how fermentation status, visitor abundance, sap chemistry, and exposure conditions influence eDNA recovery from sap surfaces.

## 5 Conclusions

Using surface environmental DNA collected from natural tree sap and sap-mimicking traps, we detected sap-associated insect taxa (BLAST top-hit identity ≥98%) at two forest sites, Fujisawa and Minamisanriku. Across sites, eDNA recovered multiple sap-associated taxa from several insect orders, including taxa that were not recorded by conventional visual monitoring during the same season. These results support the potential complementarity of sap-surface eDNA with established field surveys.

However, this study should be interpreted as a pilot proof of concept. The study included limited replication and did not include dedicated field-blank swabs, extraction blanks, or PCR negative controls; therefore, the findings should be interpreted as evidence of feasibility rather than as a demonstration of zero contamination. In addition, sap-surface eDNA cannot yet be attributed exclusively to direct sap visitation rather than environmental deposition. Differences between markers and between natural sap and sap-mimicking trap samples likely reflect a combination of primer bias, differential DNA degradation, variation in DNA shedding, and stochastic sampling of low-template DNA.

Together, our results support the feasibility of tree sap–associated surface eDNA as a monitoring resource for sap-associated insects and suggest that it may be useful for detecting taxa that are difficult to observe during conventional surveys. The contrast between active fermented sap and inactive non-fermented sap further suggests that careful selection of insect-active sap patches is important for reliable detection, although this result remains preliminary because it was based on an unreplicated spot comparison. Future work should incorporate replicated temporal sampling, dedicated process controls, optimized preservation protocols, and direct comparisons between natural sap patches and standardized sap-mimicking traps.

## 6 Data availability statement

Raw sequence reads generated in this study will be deposited in the NCBI Sequence Read Archive prior to journal publication. Processed taxon tables, curated species lists, metadata, and scripts used for figure generation will be made available in Zenodo or an equivalent public repository. Data and code will be made available no later than publication of the peer-reviewed version of this article.

## 7. Acknowledgements

We would like to express our gratitude to the members of the Ichinose Laboratory and the Sato-FC Project for their cooperation in field surveys and conventional monitoring records. We also thank Mr. Hayato Suzuki for his cooperation in field surveys in Fujisawa and Minamisanriku. Mr. Suzuki also provided the drone aerial photographs shown in Figure 2.

## 9 Author contributions

HKaw conceived the study, conducted field sampling, compiled conventional monitoring data, curated and analyzed the data, prepared the figures, and wrote the initial manuscript draft. HY contributed to study design, field sampling, data interpretation, and manuscript revision. HKur contributed to interpretation of the eDNA results, methodological discussion, and critical revision of the manuscript. TI supervised the study and contributed to study design, interpretation, and manuscript revision. All authors reviewed and approved the final manuscript.

## References

Allen, M. C., Kwait, R., Vastano, A., Kisurin, A., Zoccolo, I., Jaffe, B. D., Angle, J. C., Maslo, B., & Lockwood, J. L. (2023a). Sampling environmental DNA from trees and soil to detect cryptic arboreal mammals. Scientific Reports, 13, 180. 10.1038/s41598-023-27512-8

Allen, M. C., Lockwood, J. L., Kwait, R., Vastano, A. R., Peterson, D. L., Tkacenko, L. A., Kisurin, A., Stringham, O., Angle, J., & Jaffe, B. (2023b). Using surface environmental DNA to assess arthropod biodiversity within a forested ecosystem. Environmental DNA, 5(6), 1652–1666. 10.1002/edn3.487

Andruszkiewicz Allan, E., Zhang, W. G., Lavery, A. C., & Govindarajan, A. F. (2021). Environmental DNA shedding and decay rates from diverse animal forms and thermal regimes. Environmental DNA, 3(2), 492–514. 10.1002/edn3.141

Barnes, M. A., & Turner, C. R. (2016). The ecology of environmental DNA and implications for conservation genetics. Conservation Genetics, 17(1), 1–17. 10.1007/s10592-015-0775-4

Beng, K. C., & Corlett, R. T. (2020). Applications of environmental DNA (eDNA) in ecology and conservation: Opportunities, challenges and prospects. Biodiversity and Conservation, 29(7), 2089–2121. 10.1007/s10531-020-01980-0

Benson, D. A., Cavanaugh, M., Clark, K., Karsch-Mizrachi, I., Lipman, D. J., Ostell, J., & Sayers, E. W. (2013). GenBank. Nucleic Acids Research, 41(D1), D36–D42. 10.1093/nar/gks1195

Bohmann, K., Evans, A., Gilbert, M. T. P., Carvalho, G. R., Creer, S., Knapp, M., Yu, D. W., & de Bruyn, M. (2014). Environmental DNA for wildlife biology and biodiversity monitoring. Trends in Ecology and Evolution, 29(6), 358–367. 10.1016/j.tree.2014.04.003

Bolyen, E., Rideout, J. R., Dillon, M. R., Bokulich, N. A., Abnet, C. C., Al-Ghalith, G. A., Alexander, H., Alm, E. J., Arumugam, M., Asnicar, F., Bai, Y., Bisanz, J. E., Bittinger, K., Brejnrod, A., Brislawn, C. J., Brown, C. T., Callahan, B. J., Caraballo-Rodríguez, A. M., Chase, J., … Caporaso, J. G. (2019). Reproducible, interactive, scalable and extensible microbiome data science using QIIME 2. Nature Biotechnology, 37(8), 852–857. 10.1038/s41587-019-0209-9

Callahan, B. J., McMurdie, P. J., Rosen, M. J., Han, A. W., Johnson, A. J. A., & Holmes, S. P. (2016). DADA2: High-resolution sample inference from Illumina amplicon data. Nature Methods, 13, 581–583. 10.1038/nmeth.3869

Camacho, C., Coulouris, G., Avagyan, V., Ma, N., Papadopoulos, J., Bealer, K., & Madden, T. L. (2009). BLAST+: Architecture and applications. BMC Bioinformatics, 10, 421. 10.1186/1471-2105-10-421

Cristescu, M. E. (2014). From barcoding single individuals to metabarcoding biological communities: Towards an integrative approach to the study of global biodiversity. Trends in Ecology & Evolution, 29(10), 566–571. 10.1016/j.tree.2014.08.001

Deiner, K., Bik, H. M., Mächler, E., Seymour, M., Lacoursière-Roussel, A., Altermatt, F., Creer, S., Bista, I., Lodge, D. M., de Vere, N., Pfrender, M. E., & Bernatchez, L. (2017). Environmental DNA metabarcoding: Transforming how we survey animal and plant communities. Molecular Ecology, 26(21), 5872–5895. 10.1111/mec.14350

Elbrecht, V., & Leese, F. (2015). Can DNA-Based Ecosystem Assessments Quantify Species Abundance? PLOS ONE, 10(7), e0130324. 10.1371/journal.pone.0130324

Elbrecht, V., & Leese, F. (2017). Validation and development of COI metabarcoding primers for freshwater macroinvertebrate bioassessment. Frontiers in Environmental Science, 5, 11. 10.3389/fenvs.2017.00011

Elbrecht, V., Braukmann, T. W. A., Ivanova, N. V., Prosser, S. W. J., Hajibabaei, M., Wright, M., Zakharov, E. V., Hebert, P. D. N., & Steinke, D. (2019). Validation of COI metabarcoding primers for terrestrial arthropods. PeerJ, 7, e7745. 10.7717/peerj.7745

Ficetola, G. F., Miaud, C., Pompanon, F., & Taberlet, P. (2008). Species detection using environmental DNA from water samples. Biology Letters, 4(4), 423–425. 10.1098/rsbl.2008.0118

Fukumoto, S., Ushimaru, A., & Minamoto, T. (2015). A basin-scale application of environmental DNA assessment for rare endemic species and closely related exotic species in rivers: A case study of giant salamanders in Japan. Journal of Applied Ecology, 52(2), 358–365. 10.1111/1365-2664.12392

Goldberg, C. S., Turner, C. R., Deiner, K., Klymus, K. E., Thomsen, P. F., Murphy, M. A., Spear, S. F., McKee, A., Oyler-McCance, S. J., Cornman, R. S., Laramie, M. B., Mahon, A. R., Lance, R. F., Pilliod, D. S., Strickler, K. M., Waits, L. P., Fremier, A. K., Takahara, T., Herder, J. E., & Taberlet, P. (2016). Critical considerations for the application of environmental DNA methods to detect aquatic species. Methods in Ecology and Evolution, 7(11), 1299–1307. 10.1111/2041-210X.12595

Harper, L. R., Buxton, A. S., Rees, H. C., Bruce, K., Brys, R., Halfmaerten, D., Read, D. S., Watson, H. V., Sayer, C. D., Jones, E. P., Priestley, V., Mächler, E., Múrria, C., Garcés-Pastor, S., Medupin, C., Burgess, K., Benson, G., Boonham, N., Griffiths, R. A., … Hänfling, B. (2019). Prospects and challenges of environmental DNA (eDNA) monitoring in freshwater ponds. Hydrobiologia, 826(1), 25–41. 10.1007/s10750-018-3750-5

Hassan, S., Sabreena Poczai, P., Ganai, B. A., Almalki, W. H., Gafur, A., & Sayyed, R. Z. (2022). Environmental DNA metabarcoding: A novel contrivance for documenting terrestrial biodiversity. Biology, 11(9), 1297. 10.3390/biology11091297

Jerde, C. L., Mahon, A. R., Chadderton, W. L., & Lodge, D. M. (2011). “Sight-unseen” detection of rare aquatic species using environmental DNA. Conservation Letters, 4(2), 150–157. 10.1111/j.1755-263X.2010.00158.x

Johnson, M. D., Fokar, M., Cox, R. D., & Barnes, M. A. (2021). Airborne environmental DNA metabarcoding detects more diversity, with less sampling effort, than a traditional plant community survey. BMC Ecology and Evolution, 21, 218. 10.1186/s12862-021-01947-x

Kirse, A., Bourlat, S. J., Langen, K., & Fonseca, V. G. (2021). Metabarcoding Malaise traps and soil eDNA reveals seasonal and local arthropod diversity shifts. Scientific Reports, 11, 10498. 10.1038/s41598-021-89950-6

Lodge, D. M., Turner, C. R., Jerde, C. L., Barnes, M. A., Chadderton, L., Egan, S. P., Feder, J. L., Mahon, A. R., & Pfrender, M. E. (2012). Conservation in a cup of water: Estimating biodiversity and population abundance from environmental DNA. Molecular Ecology, 21(11), 2555–2558. 10.1111/j.1365-294X.2012.05600.x

Lynggaard, C., Bertelsen, M. F., Jensen, C. V., Johnson, M. S., Frøslev, T. G., Olsen, M. T., & Bohmann, K. (2022). Airborne environmental DNA for terrestrial vertebrate community monitoring. Current Biology, 32(3), 701–707.e5. 10.1016/j.cub.2021.12.014

Macher, T. H., Schütz, R., Hörren, T., Beermann, A. J., & Leese, F. (2023). It’s raining species: Rainwash eDNA metabarcoding as a minimally invasive method to assess tree canopy invertebrate diversity. Environmental DNA, 5(1), 3–11. 10.1002/edn3.372

Magoč, T., & Salzberg, S. L. (2011). FLASH: Fast length adjustment of short reads to improve genome assemblies. Bioinformatics, 27(21), 2957–2963. 10.1093/bioinformatics/btr507

Marques, V., Milhau, T., Albouy, C., Dejean, T., Manel, S., Mouillot, D., & Juhel, J.-B. (2021). GAPeDNA: Assessing and mapping global species gaps in genetic databases for eDNA metabarcoding. Diversity and Distributions, 27(10), 1880–1892. 10.1111/ddi.13142

Marquina, D., Esparza-Salas, R., Roslin, T., & Ronquist, F. (2019). Establishing arthropod community composition using metabarcoding: Surprising inconsistencies between soil samples and preservative ethanol and homogenate from Malaise trap catches. Molecular Ecology Resources, 19(6), 1516–1530. 10.1111/1755-0998.13071

Mauvisseau, Q., Halfmaerten, D., Neyrinck, S., Burian, A., & Brys, R. (2021). Effects of preservation strategies on environmental DNA detection and quantification using ddPCR. Environmental DNA, 3(4), 815–822. 10.1002/edn3.188

Ministry of the Environment, Japan. (n.d.). Prefectural threatened species search (Ikimono Log). Biodiversity Center of Japan. Retrieved 19 February 2026, from https://ikilog.biodic.go.jp/Rdb/pref

Montgomery, G. A., Belitz, M. W., Guralnick, R. P., & Tingley, M. W. (2021). Standards and best practices for monitoring and benchmarking insects. Frontiers in Ecology and Evolution, 8, 579193. 10.3389/fevo.2020.579193

Nakamura, A., Kitching, R. L., Cao, M., Creedy, T. J., Fayle, T. M., Freiberg, M., Hewitt, C. N., Itioka, T., Koh, L. P., Ma, K., Malhi, Y., Mitchell, A., Novotny, V., Ozanne, C. M. P., Song, L., Wang, H., & Ashton, L. A. (2017). Forests and their canopies: Achievements and horizons in canopy science. Trends in Ecology & Evolution, 32(6), 438–451. 10.1016/j.tree.2017.02.020

Porter, T. M., & Hajibabaei, M. (2018). Over 2.5 million COI sequences in GenBank and growing. PLOS ONE, 13(9), e0200177. 10.1371/journal.pone.0200177

R Core Team. (2024). R: A language and environment for statistical computing. R Foundation for Statistical Computing, Vienna, Austria.

Ratnasingham, S., & Hebert, P. D. N. (2007). BOLD: The Barcode of Life Data System (http://www.barcodinglife.org). Molecular Ecology Notes, 7(3), 355–364. 10.1111/j.1471-8286.2007.01678.x

Roger, F., Ghanavi, H. R., Danielsson, N., Wahlberg, N., Löndahl, J., Pettersson, L. B., Andersson, G. K. S., Olén, N. B., & Clough, Y. (2022). Airborne environmental DNA metabarcoding for the monitoring of terrestrial insects—A proof of concept from the field. Environmental DNA, 4(4), 790–807. 10.1002/edn3.290

Ruppert, K. M., Kline, R. J., & Rahman, M. S. (2019). Past, present, and future perspectives of environmental DNA (eDNA) metabarcoding: A systematic review in methods, monitoring, and applications of global eDNA. Global Ecology and Conservation, 17, e00547. 10.1016/j.gecco.2019.e00547

Shokralla, S., Gibson, J. F., Nikbakht, H., Janzen, D. H., Hallwachs, W., & Hajibabaei, M. (2014). Next-generation DNA barcoding: Using next-generation sequencing to enhance and accelerate DNA barcode capture from single specimens. Molecular Ecology Resources, 14(5), 892–901. 10.1111/1755-0998.12236

Taberlet, P., Coissac, E., Hajibabaei, M., & Rieseberg, L. H. (2012). Environmental DNA. Molecular Ecology, 21(8), 1789–1793. 10.1111/j.1365-294X.2012.05542.x

Thomsen, P. F., Kielgast, J., Iversen, L. L., Møller, P. R., Rasmussen, M., & Willerslev, E. (2012). Detection of a diverse marine fish fauna using environmental DNA from seawater samples. PLOS ONE, 7(8), e41732. 10.1371/journal.pone.0041732

Thomsen, P. F., & Willerslev, E. (2015). Environmental DNA—An emerging tool in conservation for monitoring past and present biodiversity. Biological Conservation, 183, 4–18. 10.1016/j.biocon.2014.11.019

Thomsen, P. F., & Sigsgaard, E. E. (2019). Environmental DNA metabarcoding of wild flowers reveals diverse communities of terrestrial arthropods. Ecology and Evolution, 9(4), 1665–1679. 10.1002/ece3.4809

Ulyshen, M. D. (2011). Arthropod vertical stratification in temperate deciduous forests: Implications for conservation-oriented management. Forest Ecology and Management, 261(9), 1479–1489. 10.1016/j.foreco.2011.01.033

Weigand, H., Beermann, A. J., čiampor, F., Costa, F. O., Csabai, Z., Duarte, S., Geiger, M. F., Grabowski, M., Rimet, F., Rulik, B., Strand, M., Szucsich, N., Weigand, A. M., Willassen, E., Wyler, S. A., Bouchez, A., Borja, A., čiamporová-Zaťovičová, Z., Ferreira, S., … Ekrem, T. (2019). DNA barcode reference libraries for the monitoring of aquatic biota in Europe: Gap-analysis and recommendations for future work. Science of the Total Environment, 678, 499–524. 10.1016/j.scitotenv.2019.04.247

Wilcox, T. M., McKelvey, K. S., Young, M. K., Jane, S. F., Lowe, W. H., Whiteley, A. R., & Schwartz, M.K. (2013). Robust detection of rare species using environmental DNA: The importance of primer specificity. PLOS ONE, 8(3), e59520. 10.1371/journal.pone.0059520

Yoneya, K., Ushio, M., & Miki, T. (2023). Non-destructive collection and metabarcoding of arthropod environmental DNA remained on a terrestrial plant. Scientific Reports, 13, 7125. 10.1038/s41598-023-32862-4

Yoshimoto, J., Kakutani, T., & Nishida, T. (2005). Influence of resource abundance on the structure of the insect community attracted to fermented tree sap. Ecological Research, 20(4), 405–414. 10.1007/s11284-005-0054-9

Young, M. R., & Hebert, P. D. N. (2022). Unearthing soil arthropod diversity through DNA metabarcoding. PeerJ, 10, e12845. 10.7717/peerj.12845

Zinger, L., Bonin, A., Alsos, I. G., Bálint, M., Bik, H., Boyer, F., Chariton, A. A., Creer, S., Coissac, E., Deagle, B. E., De Barba, M., Dickie, I. A., Dumbrell, A. J., Ficetola, G. F., Fierer, N., Fumagalli, L., Gilbert, M. T. P., Jarman, S., Jumpponen, A., … Taberlet, P. (2019). DNA metabarcoding—Need for robust experimental designs to draw sound ecological conclusions. Molecular Ecology, 28(8), 1857–1862. 10.1111/mec.15060

